# Endocardial/endothelial angiocrines regulate cardiomyocyte development and maturation and induce features of ventricular non-compaction

**DOI:** 10.1101/2020.07.25.220301

**Authors:** Siyeon Rhee, David T. Paik, Johnson Y. Yang, Danielle Nagelberg, Ian Williams, Lei Tian, Robert Roth, Mark Chandy, Jiyeon Ban, Nadjet Belbachir, Seokho Kim, Hao Zhang, Ragini Phansalkar, Ka Man Wong, Devin A. King, Caroline Valdez, Virginia D. Winn, Ashby J. Morrison, Joseph C. Wu, Kristy Red-Horse

**Affiliations:** Department of Biology, Stanford University School of Medicine, Stanford University, Stanford, CA; Stanford Cardiovascular Institute, Stanford University School of Medicine, Stanford University, Stanford, CA; Department of Medicine, Division of Cardiology, Stanford University School of Medicine, Stanford University, Stanford, CA; Institute for Stem Cell Biology and Regenerative Medicine, Stanford University School of Medicine, Stanford University, Stanford, CA; Department of Developmental Biology, Stanford University School of Medicine, Stanford University, Stanford, CA; Department of Genetics, Stanford University School of Medicine, Stanford University, Stanford, CA; Department of Obstetrics and Gynecology, Stanford University School of Medicine, Stanford University, Stanford, CA

## Abstract

Non-compaction cardiomyopathy is a devastating genetic disease caused by insufficient consolidation of ventricular wall muscle that can result in inadequate cardiac performance. Despite being the third most common cardiomyopathy, the mechanisms underlying the disease, including the cell types involved, are poorly understood. We have previously shown that endothelial cell-specific deletion of the chromatin remodeler gene *Ino80* results in defective coronary vessel development that leads to ventricular noncompaction in embryonic mouse hearts. Here, we used single-cell RNA-sequencing to characterize endothelial and endocardial defects in *Ino80*-deficient hearts. We observed a pathological endocardial cell population in the non-compacted hearts, and identified multiple dysregulated angiocrine factors that dramatically affected cardiomyocyte behavior. We identified *Col15A1* as a coronary vessel-secreted angiocrine factor, downregulated by *Ino80*-deficiency, that functioned to promote cardiomyocyte proliferation. Furthermore, mutant endocardial and endothelial cells (ECs) upregulated expression of secreted factors, such as *Tgfbi, Igfbp3, Isg15*, and *Adm*, which decreased cardiomyocyte proliferation and increased maturation. These findings support a model where coronary ECs normally promote myocardial compaction through secreted factors, but that endocardial and ECs can secrete factors that contribute to non-compaction under pathological conditions.

## Introduction

Left ventricular non-compaction (LVNC) is a rare, but potentially devastating, disease in which cardiac systolic function becomes compromised due to incomplete compaction of the ventricular wall^1^. LVNC represents the third most prevalent form of congenital cardiomyopathy in the United States, manifesting in a myriad of cardiovascular complications such as heart failure, arrhythmias, and thromboembolism^2^.

Like many congenital cardiomyopathies, the genetic basis of LVNC is highly complex and remains elusive. Human genome wide association studies have identified common gene mutations and molecular pathways related to sarcomere and sarcolemma function (e.g., MYH7, MYBP3, TPM1, TNNT2) in LVNC patient cohorts^3,4^. However, the identified mutations are also commonly found in dilated or hypertrophic cardiomyopathy patients and are not unique to LVNC.

Due to the limited understanding of the precise etiology of the disease, there currently exists no therapeutic solutions that reverse the ventricular non-compaction phenotype^2,5^. Progress towards a better understanding of LVNC and how to treat it will rely on robust animal and human cell culture experimental models. Animal models of ventricular non-compaction suggest that the pathology arises during embryonic development due to insufficient proliferation and pre-mature maturation of cardiomyocytes. This can occur when signaling pathways such as TGF-β and Notch signaling are dysregulated^1,6^. Interestingly, a number of recent studies have found that the LVNC phenotype not only manifests in the myocardium, but it is also highly correlated with improper endocardial function and/or development of the coronary endothelium^7–10^. For instance, endocardial/endothelial-specific deletion of *Fkbp1a* leads to myocardial non-compaction via upregulation of endocardial Notch1 signaling, underlining the role of myocardium-to-endocardium paracrine signaling in controlling ventricular wall formation^8^. Independently, dysregulation of Notch signaling in endocardial cells was shown to generate LVNC^9,11^. In another study, the simultaneous presence of single nucleotide variants in three cardiac transcription factors, one of which is specific to vascular cells, elicited an LVNC-like phenotype in embryonic mouse hearts. These variants similarly recapitulated the phenotype in LVNC patient-specific induced pluripotent stem cell-derived cardiomyocytes (iPSC-CMs)^10^. These observations suggest the possibility that LVNC can arise through defects in cardiac endocardium and endothelium.

We previously reported that endothelial-specific deletion of the chromatin remodeler *Ino80* also resulted in an LVNC phenotype, where impaired coronary vessel angiogenesis coincided with noncompaction of the ventricular wall^7^. In contrast, myocardial development and compaction were unaffected when *Ino80* was deleted specifically in cardiomyocytes or epicardial-derived stromal cells. In the *Tie2Cre;Ino80^fl/fl^* embryonic hearts exhibiting the LVNC phenotype, we observed that coronary vessel sprouting was compromised, and endocardial cells are unable to transition to coronary endothelial cells (ECs) due to increased E2F-mediated gene transcription and proliferation^7^. Cell culture experiments showed that ECs supported cardiomyocyte compaction independent of blood flow or oxygenation. This suggested a role of functional coronary vessel ECs in ventricular wall formation via angiocrine signaling, i.e. through secreted factors that affect neighboring cells. We therefore hypothesize that when endocardium cannot properly transition to coronary endothelium, coronary endothelium cannot provide angiocrine factors that promote compaction, and instead, endocardium secretes those that stimulate maturation and inhibit proliferation in cardiomyocytes, leading to non-compaction of the myocardial wall. Here, we use the *Tie2Cre;Ino80^fl/fl^* mouse model to identify candidate angiocrines expressed by endocardial and ECs, during normal development and LVNC, and test the effect of these candidates on cardiomyocyte proliferation and maturation.

## Results

### Altered endocardial and cardiomyocyte transcriptomes in Ino80 mutant hearts with LVNC

To begin understanding how defective endothelial/endocardial cells may be involved in noncompaction of neighboring myocardium, we performed scRNA-seq on dissected ventricles from control and *Tie2Cre;Ino80^fl/fl^* hearts, the latter of which exhibits ventricular non-compaction (**Fig. 1A**). We collected individual ventricular cells from embryonic day E15.5 hearts and performed droplet-based scRNA-seq using the 10x Genomics platform. 576 mutant and 518 control cells were used for downstream analysis (**s. Fig. 1**). Unsupervised graph-based clustering identified 9 expected cell populations (**Fig. 1B, s. Fig. 2A & B**), which were annotated based on expression of known markers and included endocardial cells (*Npr3+*), coronary vessel ECs (*Cldn5+, Fabp4+*), smooth muscle cells/pericytes (*Pdgfrb+*), epicardium (*Apela+*), three fibroblast clusters (*Tcf21+, Col1a1+, Vcan+*), and two ventricular cardiomyocyte clusters (*Tnni1+, Ttn+*). All of these cell types were observed in ventricles from mice of both genotypes (**Fig. 1C**).

**Fig. 1.**
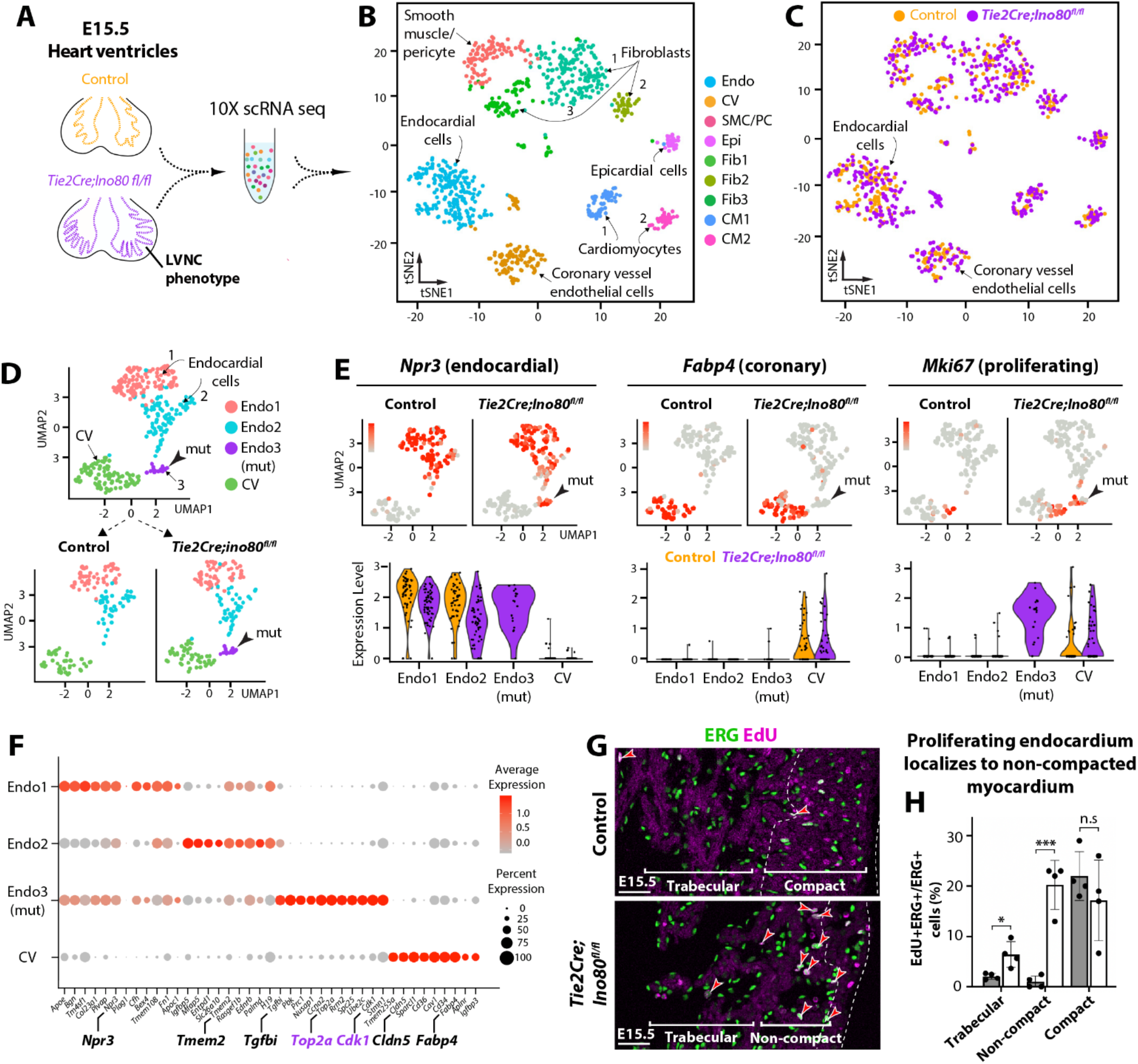
Single-cell mRNA sequencing reveals mutant-specific cell states in *Tie2Cre;Ino80^fl/fl^* heart with LVNC. **(A)** Schematic of experiment, and **(B)** t-SNE visualization of cell populations identified in the two datasets combined. **(C)** tSNE visualization of control and mutant cells. **(D)** UMAP analysis of endothelial and endocardial populations reveals a mutant-specific endocardial cell cluster (arrow, mut). **(E)** Feature and violin plots visualizing genes specific for endocardial, coronary vessel endothelial, and proliferating cells. Mutant-specific cluster represents proliferative endocardium. **(F)** Expression of top 10 genes that define each cluster. **(G)** EdU-positive endocardial cells (red arrows) are increased in mutant hearts and mostly localized to non-compacted myocardium. Control, n = 4; mutant, n = 4. **(H)** Quantification of G where each black dot represents the average of >3 FOVs from one heart. Endo: endocardial cells; CV: coronary vessel ECs; SMC: smooth muscle cells; PC: pericytes; Epi: epicardial cells; Fib: fibroblasts; CM: cardiomyocytes. Scale bars: 25 μm (G). n.s: non-significant; *P<0.05; ***P < 0.001, evaluated by Student’s t-test. Error bars are mean ± sd.

**Fig. 2.**
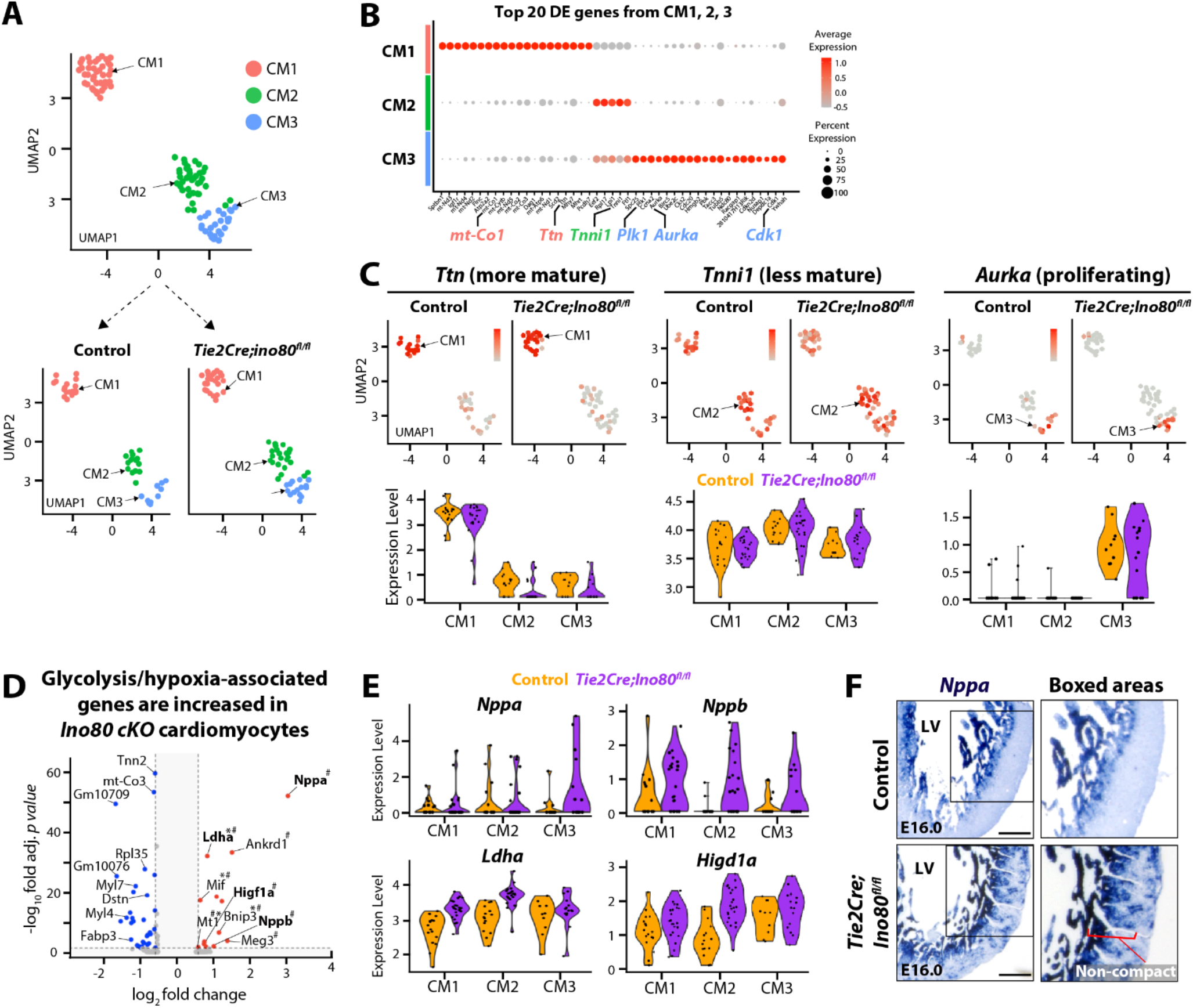
Effects of endothelial/endocardial deletion of *Ino80* on cardiomyocyte gene expression. **(A)** Subclustering of cardiomyocytes (CMs) visualized on UMAPs. **(B)** Expression of top genes that define each subcluster. Highlighted genes suggest: CM1=more mature, CM2=less mature, CM3=proliferating. **(C)** Feature and violin plots of select CM cluster-defining genes. **(D)** Volcano plot showing differentially expressed genes. Blue indicates decreased in mutant; red indicates increased in mutant where significance was considered < 0.05 adjusted p-value evaluated by Poisson method. #: hypoxia-associated genes, *: glycolysis-associated genes. **(E)** Violin plots of select genes showing how differences are spread over three CM clusters. **(F)** *In situ* hybridization validated *Nppa* expression changes. Images are representative of: control, n = 3; mutant, n = 3 hearts. Scale bars: 25 μm.

We next subclustered the endocardial and coronary endothelial populations to specifically investigate the differences between control and *Ino80* mutant vascular cells. Subclustering identified four clusters within this population—three were comprised of endocardial cells and one contained coronary endothelium (**Fig. 1D**). One of the three endocardial clusters (Endo 3) contained only *Ino80*-deficient cells and no control cells (**Fig. 1D, lower panel**). This mutant-specific cluster was defined in large part by high expression of proliferation genes, such as *Top2a* and *Cdk1*, which are very low in normal endocardium at this stage of development (**Fig. 1E & F, s. Table 1**). Consistent with previous analyses from E12.5^7^*, in vivo* EdU-labeling and tissue sectioning revealed that these ectopically cycling endocardial cells primarily localized to non-compacted myocardium at E15.5 (**Fig. 1G & H**). Thus, this mouse model of ventricular non-compaction is coincident with the emergence of mutant-specific proliferative endocardial cells embedded within non-compacted myocardium.

Since the phenotype following endothelial/endocardial *Ino80* deletion is manifested in the myocardium, we next subclustered cardiomyocytes to observe transcriptional changes associated with non-compaction. Following subclustering, we observed three prominent cardiomyocytes subtypes, all of which contained cells from control and mutant hearts (**Fig. 2A**). We first expected that the different clusters would represent compact and trabecular myocardium. However, known compact and trabecular marker genes did not distinguish the clusters. Instead, gene expression patterns indicated they were separated by maturity level and cell cycle. Specifically, cardiomyocyte cluster 1 (CM1) was defined by mature myocardial markers such as *Ttn*^1*2*^, *Myh7*^13^, and *mt-Co1^14^*, and CM2 was defined by higher expression of markers found in immature myocardium in other studies such as *Tnni1*^15^ (**Fig. 2B & C**). CM3 was composed of cycling cells that also had lower expression of many of the CM2 genes (**Fig. 2B & C**).

Genes that were differentially expressed between CMs from control and *Ino80*-deficient hearts were calculated (all CMs from each genotypes were combined to increase statistical power). 39 genes were downregulated in *Ino80*-deficient hearts and 13 were upregulated (**Fig. 2D**). Gene Set Enrichment Analysis (GSEA) to probe hallmark gene sets identified glycolysis, hypoxia, and fatty acid metabolism as being increased in mutants and gene classified as involved in development of skeletal muscle (myogenesis) to be down (**s. Table 2**). Specific genes in this pathway are highlighted in **Fig. 2D** and **E**. Of the differentially expressed genes, *Nppa* was validated using in situ hybridization (**Fig. 2F**). Although one limitation of these data is the low number of cells, we concluded that gene expression analysis indicates that cardiomyocyte metabolic state and development is dysregulated.

### LVNC-associated genes are expressed in multiple cardiac cell types

Our findings in *Tie2Cre;Ino80^fl/fl^* indicate that primary defects in vascular cells can cause ventricular non-compaction in mice. To ascertain whether this is also possible in humans, we assessed the expression of genes that have variants associated with LVNC in humans in the cell types from E15.5 mouse ventricles. We found that approximately one third of the human LVNC associated genes were expressed in cardiomyocytes (**Fig. 3A**). Interestingly, about half were specific to the more immature cluster, CM1, while the others were more prominently expressed in the more mature cluster, CM2. Another third was not highly expressed in cardiomyocytes, but instead found in various stromal cell types, including endocardial cells, coronary vessel ECs, smooth muscle cells, and fibroblasts (**Fig. 3A**). Since genes with variants associated with LVNC in humans suggests that their altered function could be causal, the specific expression of these in cells other than cardiomyocytes suggests the LVNC genes might function in other cell types to induce pathology.

**Fig. 3.**
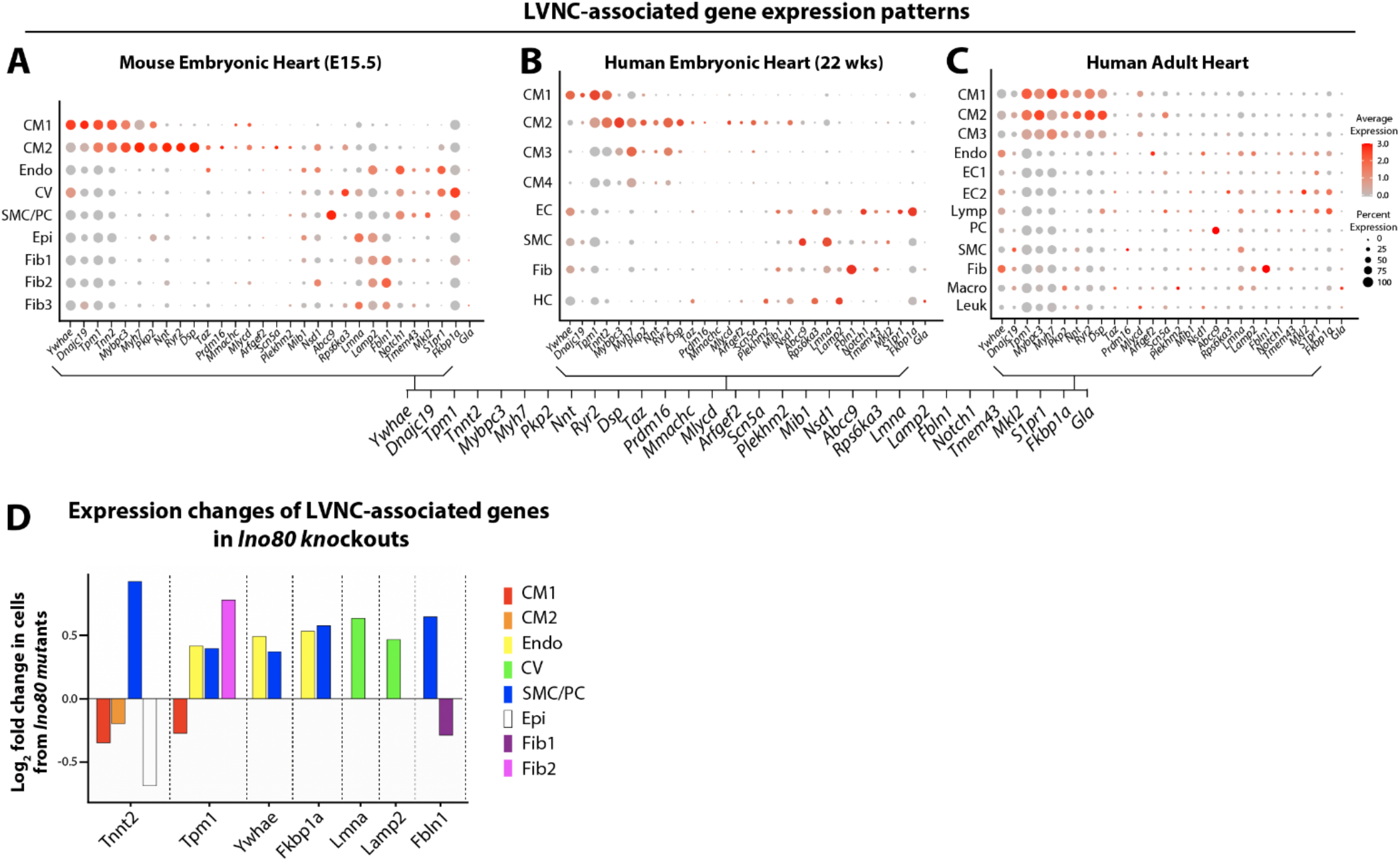
LVNC-associated genes are expressed in multiple cell types in mouse and human hearts. **(A)** Dot plots showing expression of genes with reported genetic variants associated with human LVNC in the mouse heart, and **(B-C)** in the human embryonic (22-week post gestation) and adult heart. **(D)** LVNC-associated genes that change in *Ino80*-deficient hearts. CM: cardiomyocytes, Endo: endocardial cells; CV: coronary vessel ECs; SMC: smooth muscle cells; PC: pericytes; Epi: epicardial cells; Fib: fibroblasts; HC: Hematopoietic cells; Macro: Macrophages; Leuk: Leukocytes.

To investigate the relevance of these findings to human development, we analyzed single-cell transcriptome measurements made on cells isolated from a 22-week-old human fetal heart. Expression patterns of LVNC genes in fetal human cardiac cells were similar to those observed in mice (compare **Fig. 3A & 3B**). Furthermore, analysis of a publicly available adult human heart scRNA-seq dataset^16^ revealed that the same set of LVNC-associated genes were found in both cardiomyocytes and stromal populations (**Fig. 3C**). A number of genes expressed in fetal hearts, particularly those specific to cardiomyocytes, maintained their expression in the adult heart (**Fig. 3C**). Finally, we investigated whether the genes associated with LVNC have significant expression level differences in *Ino80* mutant mice. Among these, 14 genes were differentially expressed in at least one cell type in our model of LVNC (**Fig. 3D**). These data indicate the possibility that LVNC-associated genes are derived from cells other than cardiomyocytes in humans and that they may be important for cardiac homeostasis and function in the adult. Taken together, our scRNA-seq dataset provides transcriptomic information from each cardiac cell subtypes in normal and *Ino80* mutant hearts, which we use in subsequent experiments to further understand the role of endocardial/ECs in development of ventricular non-compaction.

### Col15a1 is expressed in coronary ECs and stimulates cardiomyocyte proliferation

Our previous study showed that ECs stimulate cardiomyocyte proliferation, a behavior critical for ventricular compaction, independent of blood flow and in an *Ino80*-depedent manner^7^. We hypothesized that this activity was mediated through a factor secreted by coronary ECs that was decreased in *Ino80* mutants^7^. Our scRNA-seq data identified *Col15a1* as being significantly downregulated in mutant coronary ECs (**s. Table 3**). *Col15a1* was normally expressed in coronary ECs, smooth muscle cells, and fibroblasts in both mouse and human (**Fig. 4A**), but it was only significantly decreased in *Ino80*-deficient coronary ECs (**Fig. 4B & C**), which was validated by *in situ* hybridization on E15.5 hearts (**Fig. 4D**). These data indicate that endothelial deletion of Ino80 impairs expression of *Col15a1* in coronary vessels, which may play a role in ventricular non-compaction.

**Fig. 4.**
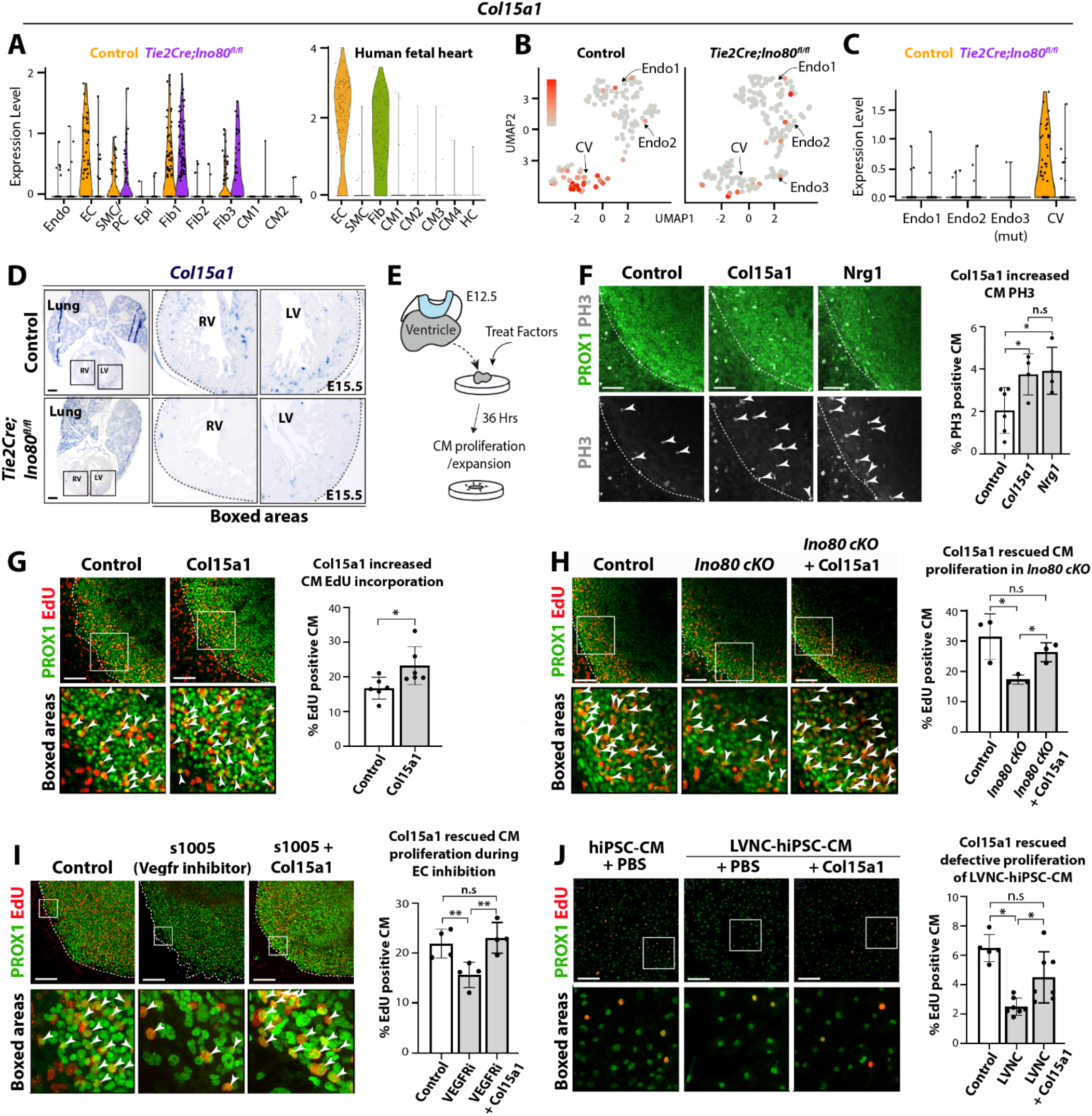
Col15a1 is decreased with LVNC and supports cardiomyocyte proliferation. **(A-C)** Violin and feature plots showing *Col15a1* expression. **(D)** *In situ* hybridization validates *Col15a1* decreased expression. Control, n = 4 hearts; mutant, n = 4 hearts. **(E)** Schematic outlining ventricle explant culture. **(F-I)** Confocal images and quantification of Col15a1-treated ventricle explants assayed for PH3 **(F)** and EdU incorporation **(G-I)**. Col15a1 increased proliferation in PROX1+ cardiomyocytes (arrowheads) **(F, G)**, which rescued the proliferative defects with *Ino80* knockout **(H)**, and with EC depletion **(I)**. **(J)** Col15a1 reversed defective proliferation in LVNC patient-specific iPSC-CMs. In bar graphs in **(F-J)**, each black dot represents the average value from >3 FOVs from individual hearts. Endo: endocardial cells; CV: coronary vessel ECs; SMC: smooth muscle cells; PC: pericytes; Epi: epicardial cells; Fib: fibroblast; CM: cardiomyocytes; HC: Hematopoietic cells. VEGFRi: VEGFR inhibitor. Scale bars: 50 μm (B); 100 μm (C-J). n.s: non-significant; *P < 0.05; **P < 0.01; ***P < 0.001, evaluated by Student’s t-test. Error bars are mean ± sd.

We next utilized heart ventricle explant cultures to assess the effect of recombinant Col15a1 on cardiomyocyte proliferation. This involved dissecting E12.5 mouse heart ventricles and culturing them in 12-well plates for 1 day. Recombinant proteins or a vehicle control was then added to the culture mediate for 36 hours prior to fixation and immunostaining^7^ (**Fig. 4E**). Proliferation was assessed by either phospho-H3 immunostaining or adding EdU to the culture 30 mins before fixing explants. Cardiomyocyte nuclei were identified through Prox1 immunofluorescence, which overlaps with Nkx2.5 but provides clearer nuclear labeling^7,17^ (**s. Fig. 3**). Adding Col15a1 increased proliferation of wild type mouse cardiomyocytes within explant cultures using both phospho-H3 localization (**Fig. 4F**) and EdU incorporation (**Fig. 4G**) as a metric, which in some cases reached levels similar to that induced by the positive control Nrg1^18^ (**Fig. 4F**). Col15a1 also rescued the lack of proliferation we had previously found to be characteristic of ventricles from *Tie2Cre;Ino80^fl/fl^* mice (**Fig. 4H**) and heart explants exposed to the VEGFR inhibitor s1005, which inhibits coronary EC growth^7^ (**Fig. 4I**).

To determine whether Col15a1 also stimulates cardiomyocyte proliferation in human cells, we utilized a human iPSC model of LVNC. Our group has previously shown that iPSC-CMs from LVNC patients exhibit defective proliferative capacity in culture^6,19^. We thus investigated whether Col15a1 could rescue this phenotype. At day 15 of the differentiation, control and LVNC iPSC-CMs with 90% purity were replated at a sparse density and treated with Col15a1 for 48 hours. Proliferation was then assessed through measuring EdU incorporation into Prox1+ cells. The results showed that Col15a1 increased proliferation in LVNC cells to levels approaching those of controls (**Fig. 4J**).

Taken together, our data suggests coronary vessel-derived Col15a1 promotes myocardial proliferation, and that its decrease in *Ino80*-deficient embryonic hearts could contribute to the non-compaction phenotype.

### Defective endocardium expresses secreted factors that inhibit cardiomyocyte proliferation and stimulate maturation

Because there was a mutant-specific endocardial cell cluster found within non-compacted myocardium, we hypothesized that those cells may secrete factors that affect neighboring cardiomyocytes during compaction. To find such factors, we looked for genes which were significantly upregulated in *Ino80* mutant endocardial and ECs that encode secreted proteins. Using this approach, we identified *Tgfbi, Igfbp3, Isg15*, and *Adm* (**Fig. 5A, s. Fig. 4, s. Table 3**) as potential mutant-specific secreted factors. The increase in expression of these genes was not specific to the Endo 3 cluster, but also occurred in other endocardial clusters and coronary endothelium (**Fig. 5A**), and, in the case of *Igfbp3*, the epicardium (**s. Fig. 4**). We were particularly intrigued by the gene *Tgfbi*, or Transforming Growth Factor Beta Induced, given our finding that human LVNC can be caused by overactive TGFb signaling^6^.

**Fig. 5.**
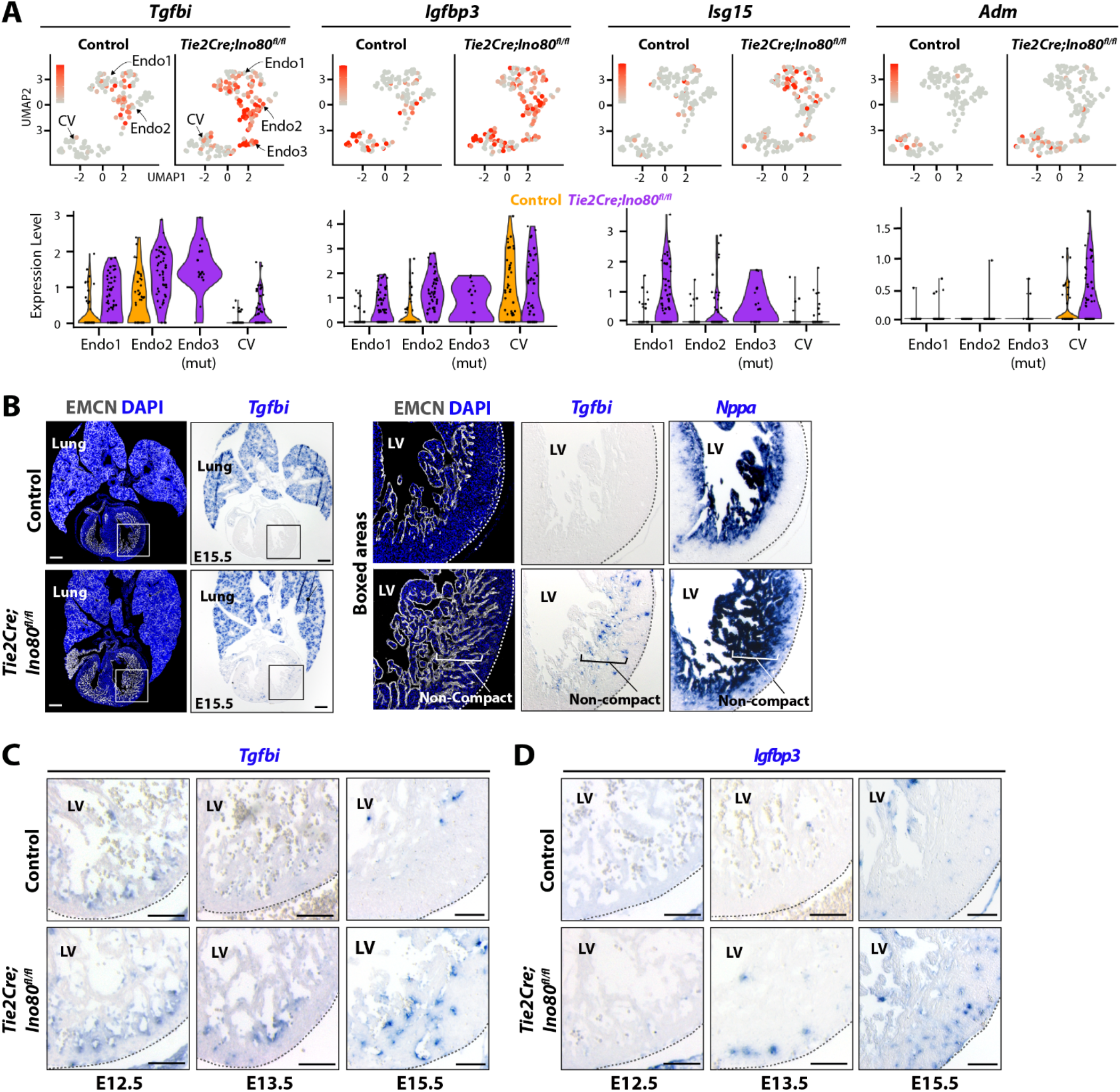
*Ino80*-deficient endocardial and ECs increase expression of secreted proteins. **(A)** Feature and violin plots of secreted factors whose expression was increased in *Ino80* mutants. **(B)** *In situ* hybridization validated *Tgfbi* expression. Expanded EMCN+ endocardium and *Nppa* expression marks non-compacted myocardium. Control, n = 4; mutant, n = 4. **(C-D)** *In situ* hybridization on left ventricles (LV) for *Tgfbi* and *Igfbp3* at indicated developmental stages. Control, n = 4 hearts; mutant, n = 4 hearts. Scale bar: 100 μm **(B)**; 25 μm **(C-D).**

We next validated these changes using *in situ* hybridization. *Tgfbi* localization in mutant and control hearts at E15.5 revealed that, while expression was similar in the lung, mutant endocardial cells within non-compacted myocardium (i.e., *Nppa+*) had increased mRNA levels (**Fig. 5B**). *In situ* hybridization at earlier stages revealed that control and mutant hearts both express *Tgfbi* at E12.5 when the heart is mostly trabeculae, although expression in mutants was qualitatively higher (**Fig. 5C**). However, most control endocardial cells downregulated *Tgfbi* by E13.5 while high expression was maintained in mutants (**Fig. 5C**). A time course of another gene increased in mutant endocardium, *Igfbp3*, showed the opposite phenomenon in that there is little endocardial expression in either genotype at E12.5, but that the gene increases to a greater extent in mutants at later time points (**Fig. 5D**). We also confirmed that the expression of *Isg15* and *Adm* were increased in mutant ventricles (**s. Fig. 5**). Thus, these genes constitute a list of secreted factors whose expression is positively correlated with ventricular non-compaction in mice.

We next aimed to address whether these gene expression changes were a direct consequence of *Ino80* deletion or perhaps a tissue level response to impaired coronary angiogenesis and/or ventricular noncompaction (e.g., hypoxia). We cross referenced the scRNA-seq gene list with bulk RNA-seq data from cultured cells and earlier staged *Tie2Cre;Ino80^fl/fl^* hearts (E13.5 & E14.5) that are not yet as severely impacted^7^. This included bulk RNA-seq performed on cultured primary human umbilical vein ECs (HUVECs) that were transfected with either control or *Ino80*-specific siRNA and whole E13.5 and E14.5 mutant hearts^7^. *Tgfbi* and *Isg15* were increased in both *Ino80* knockdown HUVECs and mutant hearts (**Fig. 6A**). However, there was either no change or a decrease in *Igfbp3* and *Adm* (**Fig. 6A**). These observations support the idea that *Isg15* and *Tgfbi* might be directly regulated by *Ino80* while *Igfbp3* and *Adm* upregulation could be a consequence of hypoxia, which is consistent with their known function of being hypoxia-inducible genes^20–22^.

**Fig. 6.**
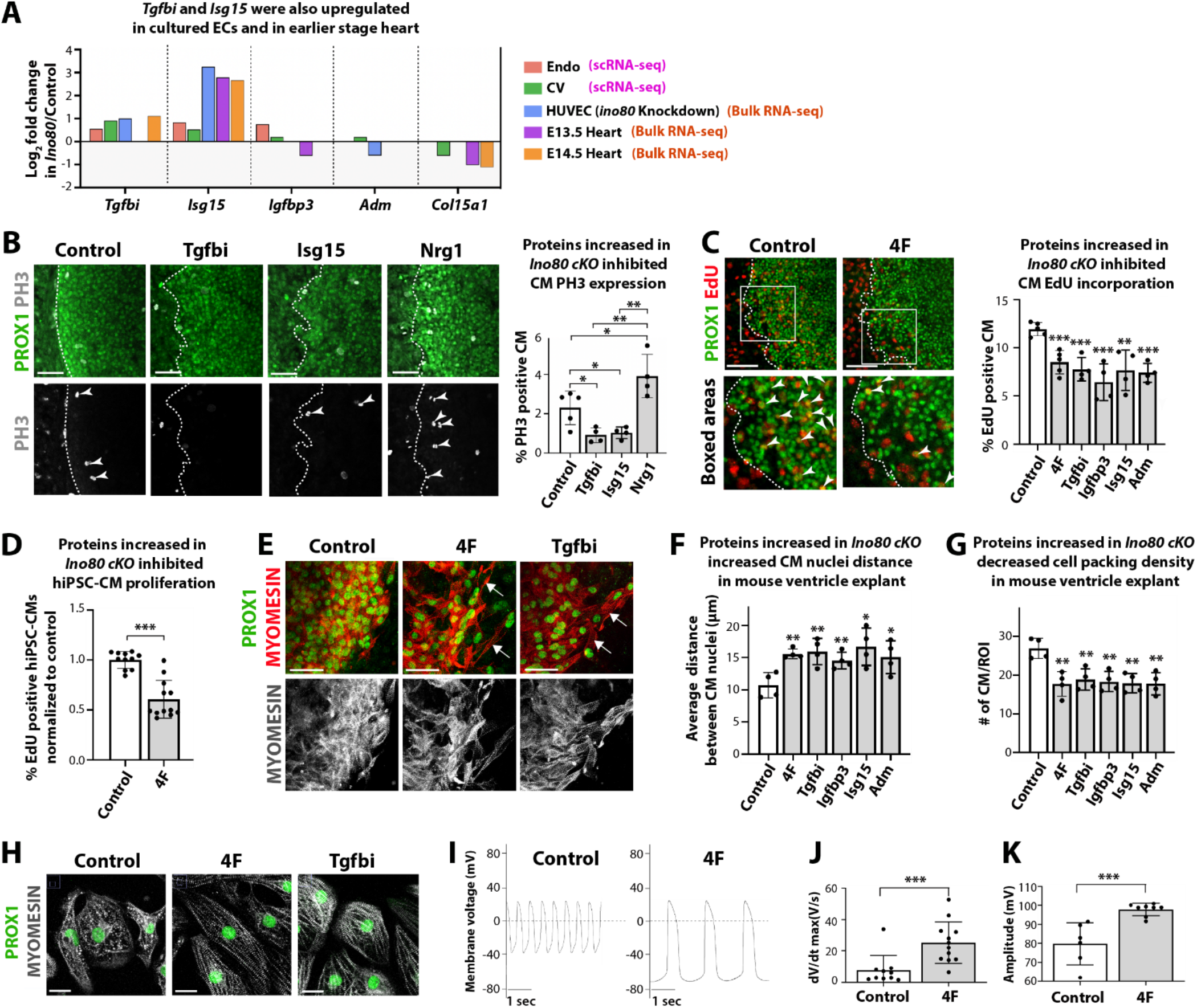
Secreted proteins upregulated in *Ino80* mutants inhibit cardiomyocyte proliferation and stimulate maturation. **(A)** Comparing scRNA-seq with bulk RNA-seq datasets from HUVECs treated with *Ino80* siRNA and whole hearts from earlier time points (E13.5 and E14.5). *Tgfbi* and *Isg15* are upregulated in cells and earlier hearts suggesting the change is not secondary to tissue hypoxia while *Igfbp3* and *Adm*, known hypoxia-inducible genes, are not. **(B-C)** Confocal images and quantification of heart ventricle explants assessed for PH3 **(B)** or EdU incorporation **(C)**. PROX1+ proliferating cardiomyocytes (arrowheads) are decreased by factors. **(D)** EdU labeling showed that treatment of 4F cocktail decreased proliferation in control iPSC-CMs. **(E)** Myomesin and PROX1 immunofluorescence revealed spreading of cardiomyocytes (arrows) in mouse ventricle explant cultures with indicated treatments. **(F)** Quantification of distance between cardiomyocytes, and **(G)** packing density of cells. **(H)** LVNC iPSC-CMs treated with 4F or Tgfbi exhibited elongated morphology with more organized sarcomeric structure. **(I)** Electrophysiological measurements of spontaneous action potentials by patch clamping of single iPSC-CMs shows 4F treatment increased maximum upstroke velocity **(J)** and action potential amplitude **(K)**. Bar graphs show N for each experiment as a black dot. Endo: endocardial cells, CV: coronary vessel ECs, HUVEC: human umbilical vein EC. Scale bars: 50 μm **(B)**; 75 μm **(C)**; 100 μm **(E, H)**. *P < 0.05; **P < 0.01; ***P < 0.001, evaluated by Student’s t-test. Error bars are mean ± sd.

We next tested the hypothesis that the secreted factors upregulated in *Ino80*-deficient endocardial and coronary ECs inhibit compaction using both mouse and human cardiomyocytes. As above, we utilized *ex vivo* assays of cardiomyocyte behavior. In addition to decreased cardiomyocyte proliferation^7^, noncompaction is associated with decreased cardiomyocyte packing (**s. Fig. 6**) and premature sarcomere maturation as indicated by increased striation development^7,23,24^. For the mouse *ex vivo* and human *in vitro* cardiomyocyte experiments, two general treatments were given, either one of the identified factors individually (*Tgfbi, Igfbp3, Isg15*, or *Adm*) or a cocktail of all factors together (4 factors or “4F”).

To investigate the exogenous effects of the four factors on cardiomyocyte proliferation, *ex vivo* mouse embryonic heart tissue cultures were prepared and analyzed as described above (**Fig. 4E**). Phospho-H3 immunostaining showed that Tgfbi and Isg15 decreased cardiomyocyte proliferation (**Fig. 6B**). EdU assays showed that cardiomyocyte proliferation was significantly decreased with application of 4F or each factor individually (**Fig. 6C**). When 4F secreted factor cocktail was treated to iPSC-CMs on day 15 of differentiation, we also observed a decrease in iPSC-CM proliferation (**Fig. 6D**). Thus, secreted factors whose expression was increased in Ino80-deficient endocardial and ECs suppress cardiomyocyte proliferation, a hallmark feature of ventricular non-compaction.

Structural changes associated with non-compaction were also assessed. It was noted that cardiomyocytes from explants treated with 4F or individual factors were less densely packed than controls (**Fig. 6E**). The average distance between nuclei has been previously used to measure cell packing^25,26^, and this analysis revealed an increase in nuclear distance and decrease in cell packing in non-compacted myocardium *in vivo* (**s. Fig. 6**). Using this metric to quantify explant cultures, we observed the same phenomenon upon application of either 4F or individual factors—they increased nuclei distance and a decreased cell packing (**Fig. 6E-G**). Therefore, these secreted factors induce a “spreading” of myocardium similar to that seen in non-compacted hearts.

Another structural change we found to be characteristic of non-compacted myocardium was premature cardiomyocyte maturation. Specifically, cardiomyocytes from *Ino80* mutant hearts exhibited more organized sarcomeres than controls at inappropriately early developmental stages^7^. Immunofluorescence for the sarcomeric protein Myomesin on 4F-treated and control iPSC-CMs showed that both 4F and Tgfbi transformed the cells from circular with punctate Myomesin to elongated with more aligned sarcomeres, which is a hallmark of maturation (**Fig. 6H**). Another feature of cardiomyocyte maturation is a change in electrophysiological properties. Action potential (AP) recordings performed by patch-clamp showed that 4F treatment increased the maximum upstroke velocity and the AP amplitude compared to the non-treated cells (**Fig. 6I-K**). Moreover, the resting membrane potential of treated cells shifted from −60 to −70 mV, a consequence of increased current through IK1 channels^27^. These changes to AP and resting membrane potential indicate an enhancement of electrophysiological maturity. Taken together, these data show that, in culture, endocardial-derived factors overexpressed in *Ino80*-mutant ECs can drive the structural phenotypes seen in non-compacted hearts.

### General inhibition of coronary angiogenesis recapitulates non-compaction

Our observation that coronary vessels express a gene for a secreted factor that stimulates cardiomyocyte proliferation, and that factors that inhibit proliferation are hypoxia inducible genes, suggests that general coronary angiogenesis could lead to non-compaction, irrespective of Ino80. To test this possibility, we specifically inhibited coronary angiogenesis using a different genetic model and analyzed ventricular wall development. *AplnCreER* was used to delete VEGFR2 at E12.5 and E13.5, which analysis performed at E16.5 (**Fig. 7A**). *AplnCreER* is specific to coronary ECs in the developing heart^28^ and VEGFR2 is required for general EC growth, allowing us to specifically inhibit coronary angiogenesis without affecting the endocardium. Quantification of trabecular and compact myocardial width in these animals at E16.5 revealed a defect in compact expansion at the expense of over-developed trabeculae (**Fig. 7B & C).** This is a hallmark of the non-compaction phenotype, and, therefore, we concluded general coronary vessel inhibition can lead to non-compaction in mice.

**Fig. 7.**
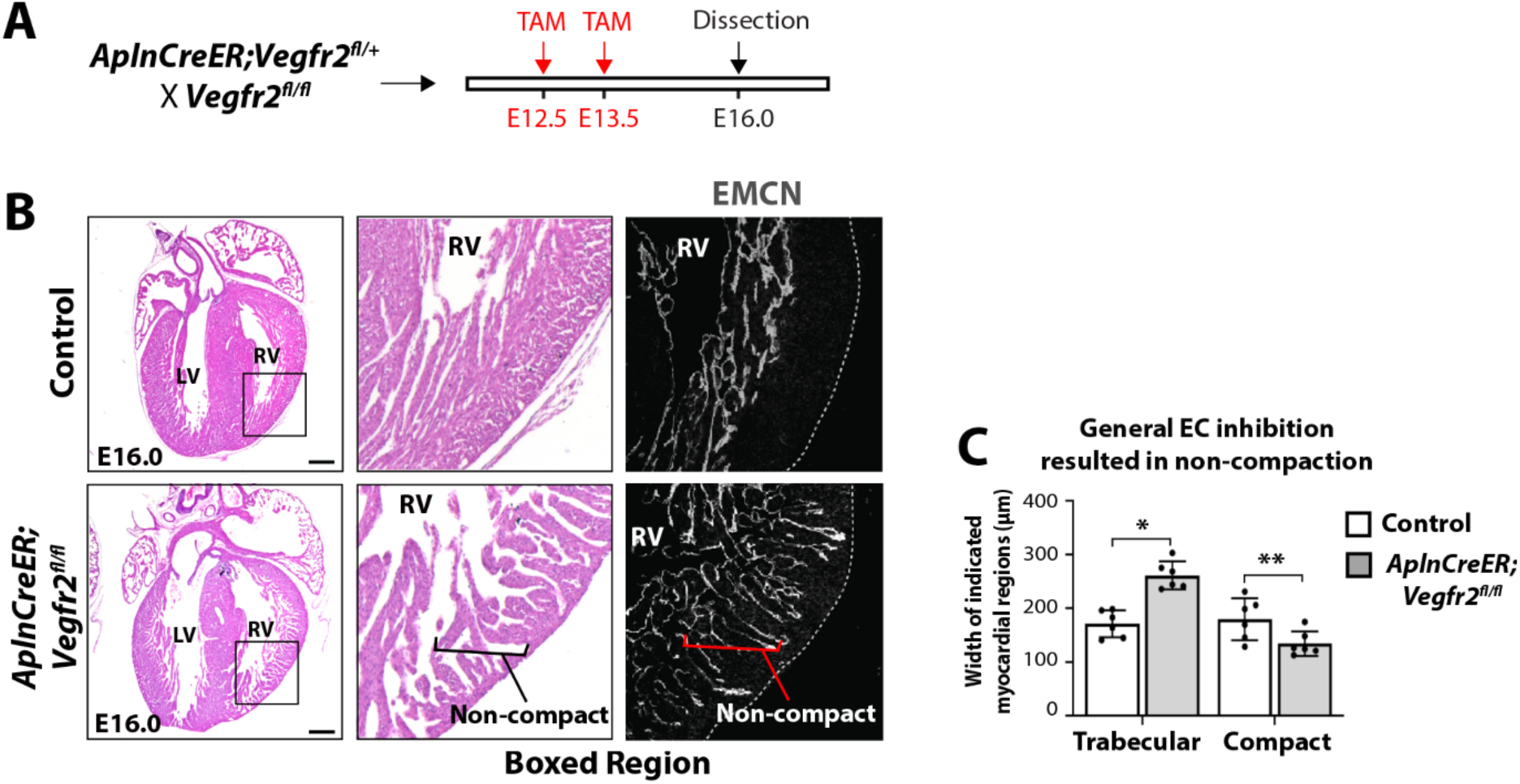
Inhibited coronary vessel growth leads to myocardial non-compaction. **(A)** Schematic showing experimental strategy to specifically inhibit coronary vessel growth. **(B)** Hematoxylin and eosin staining reveals non-compaction in *AplnCreER;Vegfr2^fl/fl^* hearts. **(C)** Quantification of compact and trabecular myocardium. Each black dot in bar graphs are the average value from >3 FOVs from one heart. Scale bars: 100 μm (B). *P < 0.05; **P < 0.01, evaluated by Student’s t-test. Error bars are mean ± sd.

### Adult phenotype of embryos exhibiting non-compaction

We next sought to understand whether the dramatic embryonic phenotype we detected with *Tie2Cre;Ino80^fl/fl^* resulted in cardiac disease in the adult. To this end, we allowed animals grow to adulthood and observed heart wall morphology and cardiac function. We first noted a profound lethality with *Tie2Cre;Ino80^fl/fl^* animals so that there were not enough adult animals of this genotype to perform a meaningful analysis (**Fig. 8A**). This lethality could result from the severe ventricular non-compaction observed in these animals, but since this is a constitutive pan-endothelial Cre that also recombines in the hematopoietic system, we cannot exclude impacts on other vascular beds and blood cells. We instead decided to use a more specific Cre, *Nfatc1Cre*, which is specific to endocardial cells in the heart^29,30^. Endocardial cells give rise to a large subset of coronary vessels, and *Nfatc1Cre;Ino80^fl/fl^* mice exhibit evidence of non-compaction, although with much less severity^7^. *Nfatc1Cre;Ino80^fl/fl^* mice survived at expected frequencies (**Fig. 8A**), and we therefore performed histological analysis and cardiac phenotyping on adults. Basic histological analyses using H&E staining revealed that *Nfatc1Cre;Ino80^fl/fl^* hearts did not have extensive trabeculations, but almost all were larger in size than controls (**Fig. 8B**). Echocardiography revealed that cardiac systolic function was significantly impaired. The left ventricular ejection fraction was decreased and end diastolic volume and left ventricular mass were increased (**Fig. 8C & D**), which were consistent with histological findings (**Fig. 8B**). These data show that endocardial and partial coronary EC deletion of *Ino80*, which causes a mild form of embryonic ventricular non-compaction^7^, leads to dysfunction in adult hearts.

**Fig. 8.**
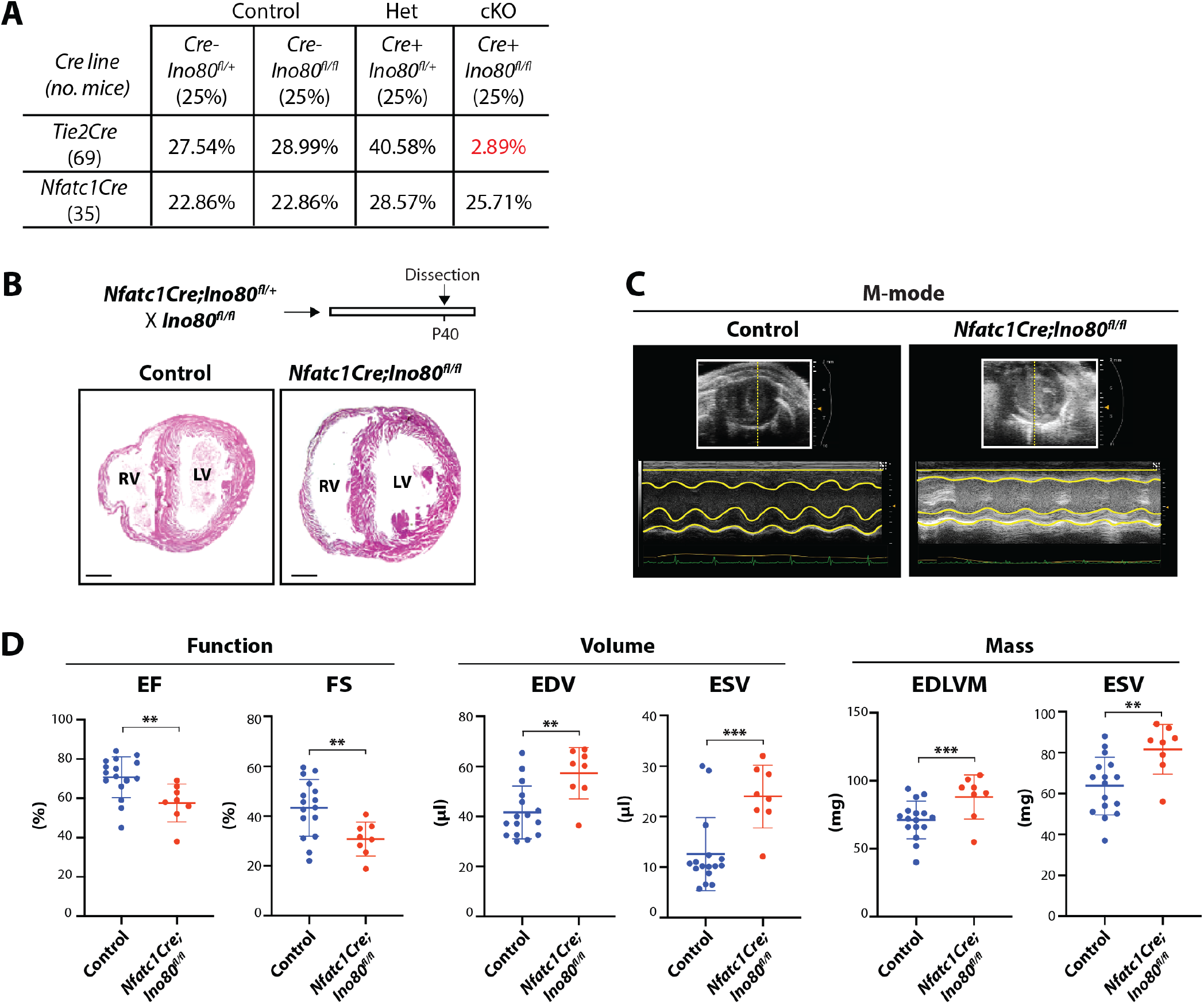
*Ino80* deletion in endocardium disrupts ventricular systolic function of adult mouse hearts. **(A)** Number of animals recovered at weaning for the indicated genotypes. **(B)** H&E staining revealed that *Nfact1Cre;Ino80^fl/fl^* mice exhibit dilated heart morphology. Control, n= 3 hearts; mutant, n = 3 hearts at P40. **(C)** M mode images acquired at the mid-papillary segment of the left ventricle (LV) during echocardiography at postnatal day 40. **(D)** Left ventricular function assessment. EF: ejection fraction, FS: fractional shortening, EDV: end diastolic volume, ESV: end systolic volume, EDLVM: end diastolic left ventricular mass, ESLVM: end systolic left ventricular mass. Scale bars: 100 μm (B), **P < 0.01; ***P < 0.001, evaluated by Student’s t-test. Error bars are mean ± sd.

## Discussion

Here, we used a mouse model with endothelial genetic insufficiency (EC-specific *Ino80* deletion) to study the potential role of vascular-derived factors in normal and pathological heart wall growth. We performed scRNA-seq of *Ino80* mutant hearts to investigate the cell state changes that accompany LVNC and to precisely characterize cell type-specific changes in gene expression. This analysis revealed a new endocardial cell state specific to *Ino80* mutant hearts that was characterized by an increase in proliferation genes, which localized to non-compacted myocardium. We also used the scRNA-seq data, as well as additional human datasets, to investigate which cells express genes associated with human LVNC. The results showed that LVNC-associated genes are expressed in both cardiomyocytes and the other stromal cell types within the heart. Thus, scRNA-seq analysis revealed that perturbations in endocardial and ECs accompany ventricular non-compaction in mice and suggested potential sites of origins for the human disease^31^.

Using scRNA-seq to identify gene expression differences within specific cells types, we identified five candidate secreted factors that endocardial or coronary ECs might use to influence cardiomyocyte behavior. Among these was *Col15A1*, which was expressed by coronary vessel ECs and was decreased in *Ino80* mutants, suggesting it might function to support cardiomyocyte proliferation. In murine explant cultures, recombinant Col15a1 stimulated cardiomyocyte proliferation. Application of Col15a1 to explant cultures that mimic LVNC—*Ino80* deficient or treated with VEGFR2 inhibitor—rescued proliferation defects. It also rescued the poor proliferative response of iPSC-CMs derived from a patient with LVNC. Little is known about the molecular function of Col15a1, but future experiments will delve into precisely how Col15a1 stimulates cardiomyocyte proliferation.

Comparing differential expression also identified several genes highly upregulated in Ino80-deficient endocardial cells and coronary ECs. This caused us to consider the hypothesis of a disease-promoting function. This is a reasonable role for the endocardium since it was overrepresented in Ino80 mutants where it lined non-compacted myocardium. Indeed, even in normal hearts, endocardial cells are associated with non-proliferative trabecular myocardium. The proteins encoded by the upregulated genes—Tgfbi, Igfbp3, Isg15 and Adm—reduced proliferation of mouse and human cardiomyocytes and induced phenotypic changes, such as elongated morphology with increased sarcomere formation and physiological maturation, consistent with it facilitating the onset of the LVNC phenotype. It will be important to explore their precise role in regulating cardiomyocyte behavior in compacting mouse hearts and in iPSC-CMs as a future direction.

Consistent with the hypoxic hallmark in cardiomyocytes from Ino80 mutant hearts, two of the genes upregulated in *Ino80* mutant endocardium and/or endothelium (*Igfbp3* and *Adm*) are known to be hypoxia-inducible and their change was not seen in non-hypoxic *Ino80* knockdown cell culture experiments or in bulk RNA-seq from stages that should be prior to onset of hypoxia. Their hypoxia inducibility along with their effects on cardiomyocytes suggests the possibility that an initial hypoxic state might start a pathologic feedback loop that exacerbates non-compaction.

In summary, our study identified and validated that cardiac endothelial- and endocardial-expressed genes encode secreted factors that can regulate cardiomyocyte proliferation and maturation. It will be important to determine whether those factors participate in organ repair or whether they are useful culture media additives that can enhance maturation of iPSC-CMs.

## Acknowledgements

This work was supported by the use of imaging instruments and image analysis support in the Stanford Center for Innovation in *In vivo* Imaging (SCi3) service center at Stanford University.

## Methods

### Mouse lines

Stanford complies with all federal and state regulations governing the humane care and use of laboratory animals, including the USDA Animal Welfare Act and our Assurance of Compliance with PHS Policy on Humane Care and Use of Laboratory Animals. The laboratory animal care program at Stanford is accredited by the Association for the Assessment and Accreditation of Laboratory Animal Care International (AAALAC Int’l). Experiments were conducted with NIH and Stanford University policies governing animal use and welfare. The following strains used were: wild-type (CD1 and FVB, Charles River Laboratories), *Tie2Cre* (The Jackson Laboratories Stock number 004128), B6.CgTg(Tekcre)12Flv/J), *Nfatc1Cre*, and *Ino80^flox^*. For experiments, Cre-expressing *Ino80^flox/+^* male mice were crossed with *Ino80^fl/fl^* females. Transgenic mice were on mixed backgrounds and ages (embryonic and adult) are indicated with each experiment. Both male and female embryos (1:1 ratio) were included in analyses as we did not genotype for gender. When using the *AplnCreER* to excise the Ino80, tamoxifen (4-OHT, H6278 Sigma) was dissolved in corn oil at a concentration of 20 mg/ml and injected into pregnant females by oral gavage at E12.5 and E13.5.

### Embryo dissection and tissue digestion for scRNA-seq

From breeding of male *Tie2Cre;Ino80^fl/+^* and female *Ino80^fl/fl^* mice, embryos were collected. Mouse ventricular heart tissue were obtained at E15.5, from which single cells were isolated and subjected to scRNA-seq. Embryos were dissected and placed in cold, sterile PBS (Life Technologies, CAT#14190250). Tail DNA was extracted and used for genotyping to distinguish control and *Ino80* cKO embryos before further microdissection. Ventricles of three hearts were micro-dissected and pooled into a 600 μl mix consisting of 1.2 U/ml dispase (Worthington #LS02100), 500 U/ml collagenase IV (Worthington #LS004186), 32 U/ml DNase I (Worthington #LS002007), and sterile DPBS with Mg^2+^ and Ca^2+^. The pooled tissue were then minced with razor blades (VWR 55411-050) and re-suspended in another 300 μl of the aforementioned mix. The pooled ventricles were then incubated at 37 °C, and gently re-suspended every 6 min. After the incubation, 60 μl cold FBS followed by 1,200 μl cold sterile PBS were added and mixed into each tube. The samples were then filtered through a 70-μm cell strainer; the filter and the source tube were washed with a total of 1,200 μl sterile PBS. Cells were then centrifuged at 400g at 4 °C for 5 min. Each cell pellet was then gently resuspended in 600 μl 3% FBS (in sterile PBS). Cells were centrifuged again at 400g at 4 °C for 5 min. Each pellet was then gently resuspended in 2,000 μl 3% FBS and 32 U/ml DNase I in sterile PBS. Cells were kept on ice until they were subjected to FACS and Clacein AM (1:1,000) stained live cells were sorted by FACS, then the sorted cells were unbiasedly subjected to single cell capture for droplet-based scRNA-seq.

### Human fetal heart tissue for scRNA-seq

Discarded human fetal left ventricle tissue (22-week-old) were collected by the Department of Obsteterics and Gynecology at Stanford University School of Medicine. The use of the discarded tissue falls under Category 1 of the OBGYN Departmental Policy on tissue donation approved by Stanford IRB. Collected tissue were minced in PBS on ice, and digested with 10 U Liberase TM enzyme (Roche 5401119001) in HBSS for 30 min at 37°C. Cells were triturated and filtered through a 100 μm (Falcon 352360) strainer and stained with 1:1,000 Calcein AM. Live cells were sorted by FACS, then the sorted cells were unbiasedly subjected to single cell capture for droplet-based scRNA-seq.

### Single-cell transcriptome library preparation and sequencing

Libraries were prepared according to the manufacturer’s instructions using the Chromium Single Cell 3’ Library & Gel Bead Kit v2 (PN-120237) and Chromium i7 Multiplex Kit (PN-120262). Final libraries were sequenced on NextSeq 500 (Illumina) by the Stanford Functional Genomics Facility. Sequencing parameters were selected according to the Chromium Single Cell v2 specifications. All libraries were sequenced to a mean read depth of at least 50,000 total aligned reads per cell. Raw sequencing reads were processed using the Cell Ranger v2.0 pipeline from 10x Genomics. Reads were demultiplexed, aligned to the mouse mm10 genome and UMI counts were quantified per gene per cell to generate a gene-barcode matrix.

### Cell filtering and cell-type clustering analysis

We sequenced the transcriptomes of approximately 500 cells captured from the control and 500 cells captured from the *Tie2Cre;Ino80^fl/fl^* embryos. Analysis of these cells were performed using the Seurat v3.1 R package^32^. Cells were excluded from downstream analysis if they expressed a) fewer than 2000 genes, b) more than 6500 genes, or c) if more than 10% of reads were aligned to mitochondrial genes. Raw UMI counts were first normalized to library size and log-transformed. We then integrated control and mutant datasets using the approach described by Butler *et al*^32^. Following integration, we reduced the dimensionality of the dataset by identifying highly variable genes and performing principal component analysis. The top 50 principal components (PCs) were used for clustering and visualization. Unsupervised clustering was performed by embedding cells in a shared nearest-neighbor graph (k=20) and then running the Louvain community detection algorithm (resolution = 0.8). Cells were projected into 2-dimensional space for visualization with the Uniform Manifold Approximation and Projection (UMAP) algorithm^33^. To identify cluster markers and differentially-expressed genes between genotypes, clusters were compared pairwise using the Wilcoxon rank sum test with a Bonferroni correction for multiple comparisons. Following cluster annotation, we subclustered and analyzed individual cell types (i.e., endocardium/endothelium, myocardium, fibroblasts, from **Fig. 1** and the endocardium reclustering discussed in **Fig. 1**). The number of PCs used and clustering resolution parameter were adjusted for these smaller subclusters.

### Histology and *in situ* hybridization

Embryos from timed pregnancies (morning of plug designated E0.5) were isolated in PBS and then fix overnight in 4% paraformaldehyde (PFA) at 4 °C. The following day, embryos were washed in PBS and PBT (PBS containing 0.1% Tween-20), dehydrated in an ascending methanol sequence, xylene treated, embedded in paraffin, and sectioned at 7 μm. Hematoxylin and eosin (H&E) staining was performed on deparaffinized slides as reported^34^. *In situ* hybridization on paraffin section was performed as described previously^9^. Briefly, dissected embryos were fixed in 4% PFA in diethyl pyrocarbonate (DEPC)-treated PBS at 4 °C overnight, washed in PBT, dehydrated in methanol series, cleared in xylene and embedded in paraffin. Embryos were sectioned at 7 um thickness. Slides were pre-incubated in hybridization buffer at 65 °C for 2 h then incubated with 1 μg/ml probes in hybridization buffer at 65 °C overnight. Antisense Tgfbi, Igfbp3, Isg15, Adm, Col15a1, Nppa probes were labeled with digoxigenin (DIG)-UTP using the Roche DIG RNA labeling System according to the manufacturer’s guidelines. After washing in salt sodium citrate (SSC) buffer, slides were incubated with alkaline phosphatase-conjugated antidigoxigenin-alkaline phosphatase antibody (Roche, 11093274910) at RT for 2 h and BM purple alkaline phosphatase substrate (Sigma-Aldrich, 1144207001) was used to visualize the color in the humid chamber. Slides were mounted using entellan mountain medium (Electron Microscopy Sciences, 14800) and imaged using Zeiss Axioimager A2 upright microscope.

### Immunohistochemistry

Immunofluorescence was performed on 7 μm deparaffinized sections^34^. Briefly, sections were subjected to antigen retrieval in Tris buffer pH 10.0 for 10 min, washed in 0.1% PBT and incubated in blocking buffer (0.5% dried milk powder, 99.5% PBT) for 2 h at room temperature (RT). Primary antibodies were incubated in blocking buffer overnight at 4 °C. The following day, the sections were washed three times with PBT and incubated for 1 h with corresponding secondary antibodies in blocking buffer at RT. After three washes in PBS, DAPI (SigmaAldrich, 1:2000) was added to counter-stain the nuclei. The sections were mounted using Prolong Gold Antifade Reagent (Invitrogen, P36934) and imaged using Zeiss LSM-700 confocal microscope. The following primary antibodies were used: ERG (Abcam, ab92513, 1:1000), ENDOMUCIN (Santa Cruz, sc-65495, 1:250), hPROX1 (R&D Systems, AF2727, 1:250), NKX2.5 (Santa Cruz, sc-8697, 1:250). Secondary antibodies were Alexa Fluor conjugates 488, 555, and 647 (Life Technologies) at 1:500.

### Explant culture and immunostaining

Whole ventricles were dissected from E12.5 embryos. Samples were rinsed with cold PBS to remove blood cells and placed in the Matrigel (1:200, BD Biosciences) with culture media (EGM-2 MV, Clonetics, CC-4147) in 24-well plates (Costar, 3524) or Cell culture inserts (Millicell, PI8P01250). Explants were cultured in 5% O2, 5% CO2 at 37 °C for 36 h before samples were fixed with 4% PFA in PBS for 15 min. After fixation, explants were washed three times with 0.5% PBT and immunofluorescence staining was performed directly within the 24-well culture plates. Explants were incubated with primary antibodies in 0.5% PBT overnight at 4 °C. Explants were washed with 0.5% PBT six times for 6 h and then incubated with corresponding secondary antibodies in 0.5% PBT overnight at 4 °C. The day after, explants were washed three times and nuclei were counter-stained with DAPI. After three washes in PBS, explants were placed in PBS and imaged using an inverted Zeiss LSM-700 confocal microscope. Captured images were digitally processed using ImageJ (NIH) and Photoshop (Adobe Systems). Antibodies used were: ERG (Abcam, ab92513, 1:1,000), Tnnt2 (DSHB, CT3-3, 1:1000), hPROX1 (R&D Systems, AF2727, 1:250). Secondary antibodies were Alexa Fluor conjugates (488, 555, 647, Life Technologies; 1:500). Smallmolecule inhibition was performed by the addition of Axitinib (Selleckchem, s1005, 20 μM) dissolved in DMSO directly to culture media after one day culture. An equal concentration of DMSO was added to the control media. To quantify proliferation of cardiomyocytes, explants were immunostained with EdU, PH3, hPROX1 (cardiomyocytes) and Myomesin (cardiomyocytes).

### Quantification of % phospho-H3 positive cardiomyocyte in ventricle explant

For cultured ventricle explants, experiment was performed three times. For each of the n=6 control hearts, the n=4 Col15a1 treated hearts and the n=4 Nrg1 treated hearts, images of the whole mount explant were taken using a Zeiss LSM-700 and maximum projection images of confocal z stacks of right or left lateral side was quantified. Proliferating cardiomyocytes were blindly counted from whole mount images as the percentage of PH3+ PROX1+ cells in a 100 μm^2^ field of view (FOV) using the Cell Counter plugin in the ImageJ (NIH) software. Three-Five FOVs were taken along the lateral sides of the heart. Unpaired two tailed t test was used to calculate P value (Prism 8.4, GraphPad; Excel 2016, Microsoft). For each of the n=6 control explant heart, the n=4 Tgfbi treated hearts, the n=4 Isg15 treated hearts, the n=4 Nrg1, PH3 positive cardiomycytes were quantified in a same way.

### Human iPSC-derived cardiomyocyte culture

Human iPSCs from one LVNC patient and healthy donors were obtained from the Stanford Cardiovascular Institute iPSC Biobank. iPSCs were cultured in E8 media (Gibco) for maintenance of pluripotency state. At 95% confluency, iPSCs were subjected to chemically defined cardiomyocyte differentiation protocol as previously described^6^. In brief, iPSCs were treated with 6 μM CHIR99021 (Selleckchem) in RPMI+B27 without insulin (Gibco) from day 0 to 2. Media was changed to RPMI+B27 without insulin from day 2 to 3, then the cells were treated with 5 μM IWR (Selleckchem) in RPMI+B27 without insulin from day 3 to 5. Media was then changed to RPMI+B27 without insulin from day 5 to 7. Robust beating of cardiomyocytes was observed starting at day 8 of differentiation. From day 7 and forward, iPSC-CMs were cultured in RPMI+B27 with insulin (Gibco). Glucose starvation from day 11 to 13 was used to eliminate non-cardiomyocyte cells. At day 15, iPSC-CMs were transferred to Matrigel-coated chamber slides or 24-well plates at a sparse cell density and cultured at 5% O^2^, 5% CO^2^ at 37 °C for 2 days. Preparation for immunostaining was performed as described above.

### Quantification of compact myocardium thickness and trabecular length

To visualize the structure of ventricles, immunostaining was performed on paraffin sections with anti-Endomucin for endocardial cells, anti-ERG for ECs and anti-hPROX1 for cardiomyocytes. ImageJ software was used to measure the thickness of the compact myocardium and the length of trabecular myocardium in tissue sections from equivalent coronal planes of the heart. The compact and trabecular myocardium regions were defined by morphology, endocardial marker staining in the heart. For each parameter, six measurements were taken along the lateral sides of the heart and averaged individually for both the left and right ventricle.

### Quantitation of distance between nuclei and cell packing density

The distance between cardiomyocytes nuclei were measured *in vivo* and explant culture using images taken with confocal microscope. Maximum projections of multiple z-stacks were created. Using measurement tool in the ImageJ 2.0.0 (NIH) software, length of distance between PROX1+ Myomesin+ cells were measured in control and *Tie2Cre;ino80^fl/fl^* heart.

### Proliferation assays

Cell proliferation *in vivo* and in explants was quantified by detection of EdU incorporation. For *in vivo* proliferation rate, 50 μg/g of body weight of EdU was injected into pregnant mice intraperitoneally 3 h before embryo collection. Proliferating ECs were calculated from paraffin tissue sections as the percentage of ERG-positive ECs labeled EdU in a 100 μm^2^ field of view. For explant cultures, 200 ng of EdU was added directly into 0.5 ml of media 30 min before fixing tissues. EdU-positive cells were detected with the Click-iT EdU kit (Invitrogen, C10338) according to manufacturer’s instruction. Briefly, Click-iT reaction cocktails were incubated for 30 min after the secondary antibody incubation of the immunostaining protocol (see above). For E15.5, experiment was performed four times. For each of the n=5 control hearts and the n=4 *Tie2Cre;ino80^fl/fl^* hearts, maximum projection images of confocal z stacks was chosen from trabecular, non-compact and compact myocardium area. Proliferating endocardial/ECs were calculated from paraffin tissue sections as the percentage of ERG positive ECs labeled EdU in a 100 μm^2^ field of view. Three-Five FOVs were taken along the lateral sides of the heart. ECs in trabecular, non-compact and compact myocardium area were identified by endothelial marker staining and anatomical locations in the heart. The total number of ECs and the number of cells positive for EdU were counted blindly using the Cell Counter plugin in the ImageJ 2.0.0 (NIH) software, the number of cells EdU positive cells was divided by total number of ECs in each regions. A two-tailed unpaired *t* test was used to determine *P* value.

For each of the n=6 control hearts and the n=6 Col15a1 treated hearts, images of the whole mount explant were taken using a Zeiss LSM-700 and maximum projection images of confocal z stacks of right or left lateral side was quantified. Proliferating cardiomyocytes were blindly counted from whole mount images as the percentage of EdU+ PROX1+ cells in a 100 μm^2^ field of view (FOV) using the Cell Counter plugin in the ImageJ (NIH) software. Three-Five FOVs were taken along the lateral sides of the heart. Unpaired two tailed t test was used to calculate P value (Prism 8.4, GraphPad; Excel 2016, Microsoft).

### Echocardiography

The left ventricular (LV) function was analyzed using a Vevo 2100 system (VisualSonics) with a 30-MHz transducer (MS400). Images were acquired at greater than 200 frames/s. Mice were anesthetized with 1-2% isofluorane and placed in a supine position imaging handling platform. Heart rate was monitored via electrocardiogram (ECG) and the mouse body temperature was maintained at 37 °C with the platform heating system. M mode and B mode images were acquired in the parasternal long axis (PLA) and three orthogonal (mid-ventricular, apical and basal) parasternal short axis (PSA) orientations. Image analysis was performed using Vevo LAB (v3.1.1). The Simpson’s biplane analysis was performed as described previously^35^. Briefly, the Vevo strain software suite was used to trace the endocardium and epicardium of the LV in the PLAX orientation. The Vevo strain integrated software tool created the summation of stacked circular disks as described in Lang et al, 2005. The LV function was quantified with the ejection fraction (EF) using the formula EF = (EDV—ESV)/EDV * 100.

### Cellular electrophysiology

At day 20 of cardiac differentiation, iPSC-CMs have been plated on 35mm glass bottom petri dishes (Cellvis) coated with Matrigel (Corning) after being enzymatically dispersed using TrypLE 10X (Gibco, Thermofisher). Patch Clamp experiments have been performed 10 days after plating in perforated patch configuration using borosilicate thin wall filaments pulled with a horizontal puller (sutter instrument) to obtain electrodes of 4 MΩ of resistance. The bath solution contains mM: 140 NaCl, 4 KCl, 1 CaCl2, 1 MgCl2, 10 HEPES, 10 Glucose, the Intra-pipette solution contains mM: 5 NaCl, 20 KCl, 20 K-Gluconate, 5 HEPES, 0,85 of Amphotericin B. Action potential signals have been recorded at 37°C in current-clamp configuration and paced at 1hZ frequency using EPC-10 patch clamp amplifier (HEKA, Lambrecht, Germany) and investigated cardiomyocytes were selected using inverted microscope (Nikon, Tokyo, Japan).

### Statistical analysis

Statistical analyses were performed using Prism (GraphPad). Data are represented as mean ± sd. For animal knockout studies, no statistical methods were used to predetermine sample size; sample size was determined based on mouse genetics. Crosses were performed until a minimum of 3–10 experimental animals (i.e., mutants) from multiple litters were obtained. No randomization or blinding was performed. Litter mate controls were always used for analyzing experimental animals. Unpaired t-test (two-tailed) were performed to assess statistical significance between two sample groups. A p < 0.05 was considered statistically significant.

