## Supplemental Figures for "Endocardial/endothelial angiocrines regulate cardiomyocyte development and maturation and induce features of ventricular non-compaction"

**Rhee, Paik, et al.**

### Supplementary Figure 1

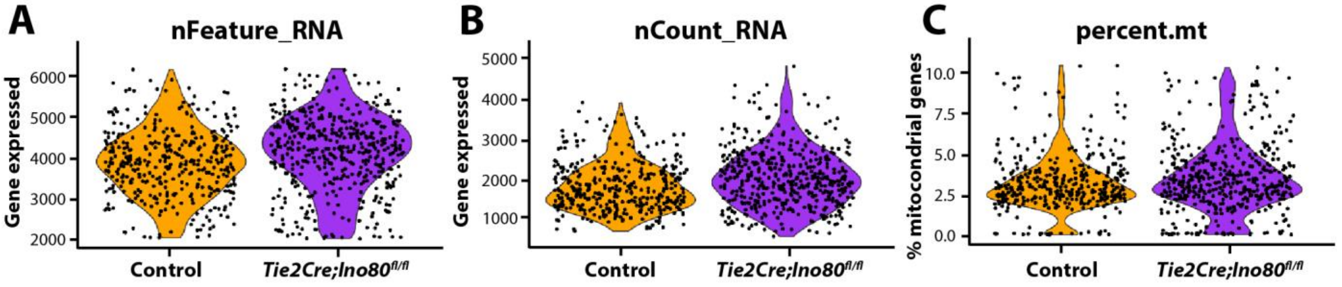


**D**

**Control**

| **Sequencing** | |
|---|---|
| Number of Reads | 152,308,822 |
| Valid Barcodes | 98.40% |
| Sequecning Saturation | 91.30% |
| Q30 Bases in Barcode | 97.40% |
| Q30 Bases in RNA Read | 84.40% |
| Q30 Bases in Sample Index | 94.60% |
| Q30 Bases in UMI | 97.20% |

| **Mapping** | |
|---|---|
| Reads Mapped Confidently to Transcriptome | 67.10% |
| Reads Mapped Confidently to Exonic Regions | 69.50% |
| Reads Mapped Confidently to Intronic Regions | 14.70% |
| Reads Mapped Confidently to Intergenic Regions | 3.40% |
| Reads Mapped Antisense to Gene | 3.60% |

| **Cells** | |
|---|---|
| Estimated Number of Cells | 518 |
| Fraction Reads in Cells | 94.30% |
| Mean Reads per cell | 294,032 |
| Median Genes per Cell | 3,584 |
| Total Genes Detected | 16,701 |
| Median UMI Counts per Cell | 14,102 |

***Tie2Cre;Ino80*** $^{fl/fl}$

| **Sequencing** | |
|---|---|
| Number of Reads | 194,000,501 |
| Valid Barcodes | 98.40% |
| Sequecning Saturation | 91.30% |
| Q30 Bases in Barcode | 97.40% |
| Q30 Bases in RNA Read | 84.40% |
| Q30 Bases in Sample Index | 94.60% |
| Q30 Bases in UMI | 97.20% |

| **Mapping** | |
|---|---|
| Reads Mapped Confidently to Transcriptome | 68.40% |
| Reads Mapped Confidently to Exonic Regions | 70.80% |
| Reads Mapped Confidently to Intronic Regions | 13.70% |
| Reads Mapped Confidently to Intergenic Regions | 3.50% |
| Reads Mapped Antisense to Gene | 3.80% |

| **Cells** | |
|---|---|
| Estimated Number of Cells | 576 |
| Fraction Reads in Cells | 94.80% |
| Mean Reads per cell | 336,806 |
| Median Genes per Cell | 4,081 |
| Total Genes Detected | 16,841 |
| Median UMI Counts per Cell | 17,624 |

**s. Fig. 1. Quality control metrics for scRNA-seq data.** Violin plots showing **(A)** the distribution of genes per cell, **(B)** number of unique molecular identifiers per cell, and **(C)** mitochondrial gene percentage split by genotype. **(D)** Parameters of sequencing data obtained from Cell Ranger (10x Genomics).

### Supplementary Figure 2

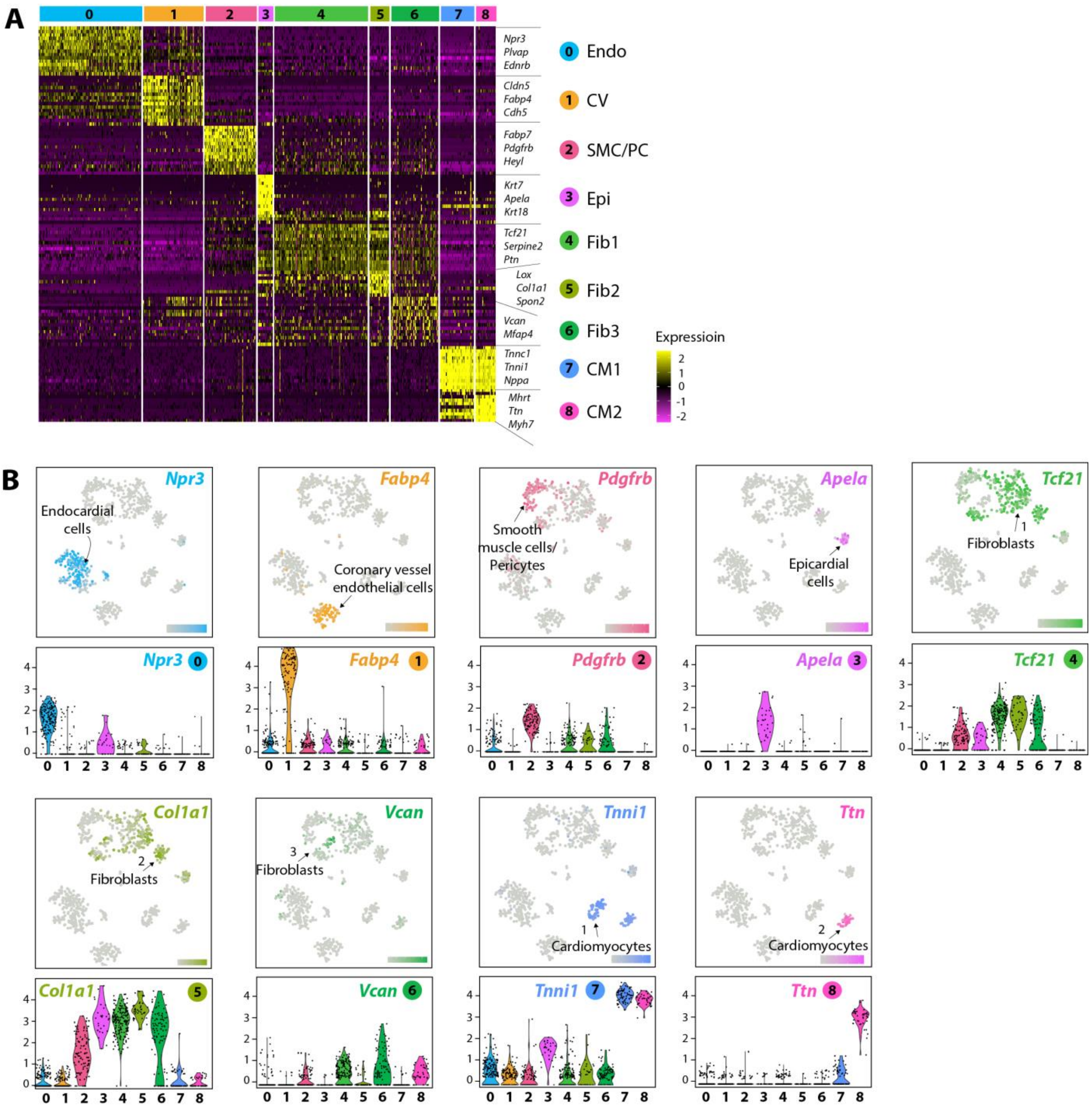


**s. Fig. 2. Identification of cell types in E15.5 mouse hearts. (A)** Heatmap showing expression of top 10 genes that define each cluster labeled in Fig. 1B. **(B)** t-SNE (top) and violin plots (bottom) of representative genes that define each cell type. Endo: endocardial cells; CV: coronary vessel endothelial cells; SMC: smooth muscle cells; PC: pericytes; Epi: epicardial cells; Fib: fibroblasts; CM: cardiomyocytes.

### Supplementary Figure 3

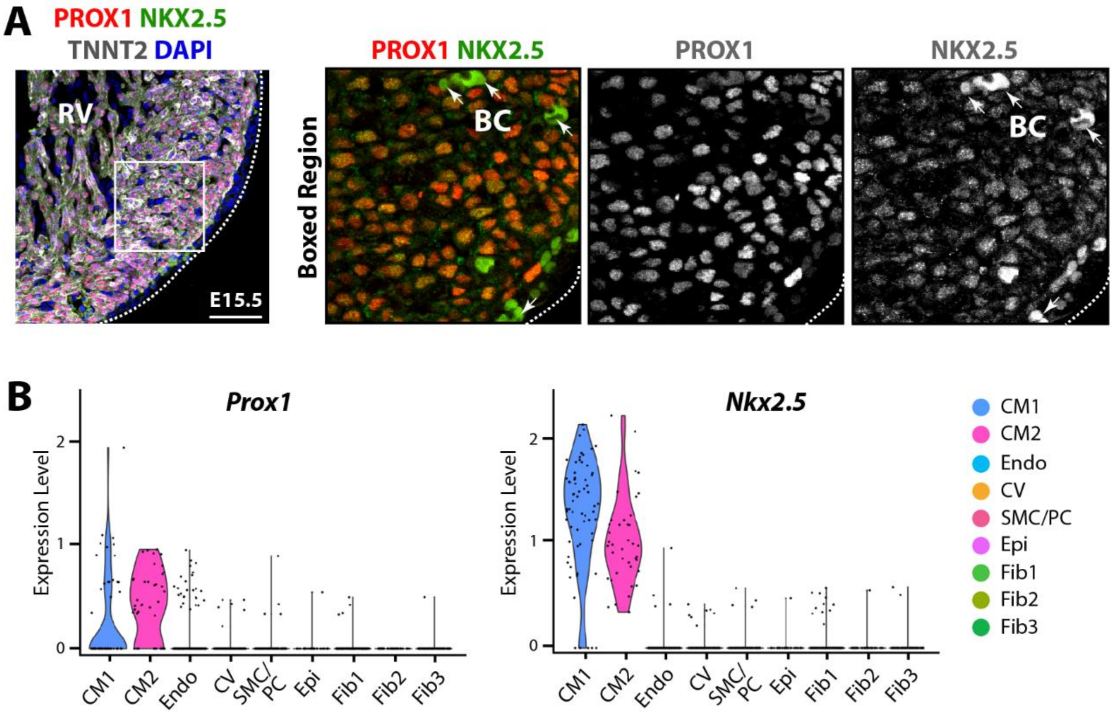


**s. Fig. 3. Prox1 is a cardiomyocyte marker in the embryonic mouse heart. (A)** Immunofluorescence for PROX1 and two well-established cardiomyocyte markers, NKX2.5 and TNNT2. PROX1 directly overlaps with NKX2.5. Images are representative of the following number of replicates: control, n = 3 hearts; mutant, n = 3 hearts at E15.5. **(B)** Violin plot shows *Prox1* is specific for cardiomyocytes in E15.5 mouse hearts. RV: right ventricle; BC: blood cells; CM: cardiomyocytes; Endo: endocardial cells; CV: coronary vessel endothelial cells; SMC: smooth muscle cells; PC: pericytes; Epi: epicardial cells; Fib: fibroblasts. Scale bars: 25 µm (A).

### Supplementary Figure 4

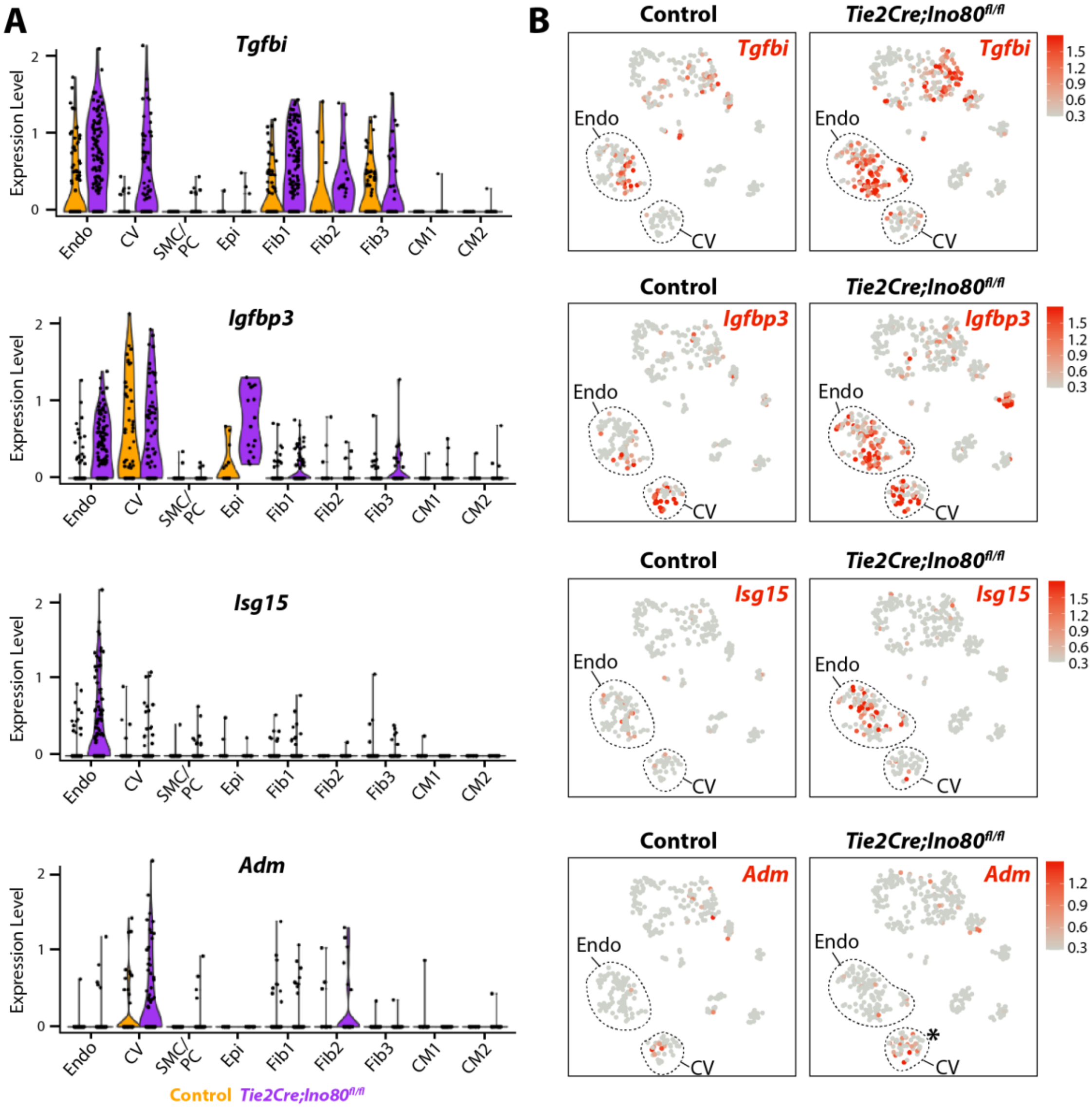


**s. Fig. 4. Expression of genes for secreted proteins in all cardiac cells in scRNA-seq dataset. (A)** Violin and **(B)** feature plots show upregulation in endocardium and/or coronary endothelial cells of all four genes. *Tgfbi*, *Igfbp3*, and *Adm* are also up-regulated in other cell populations. Endo: endocardial cells; CV: coronary vessel endothelial cells; SMC: smooth muscle cells; PC: pericytes; Epi: epicardial cells; Fib: fibroblasts; CM: cardiomyocytes.

#### Supplementary Figure 5

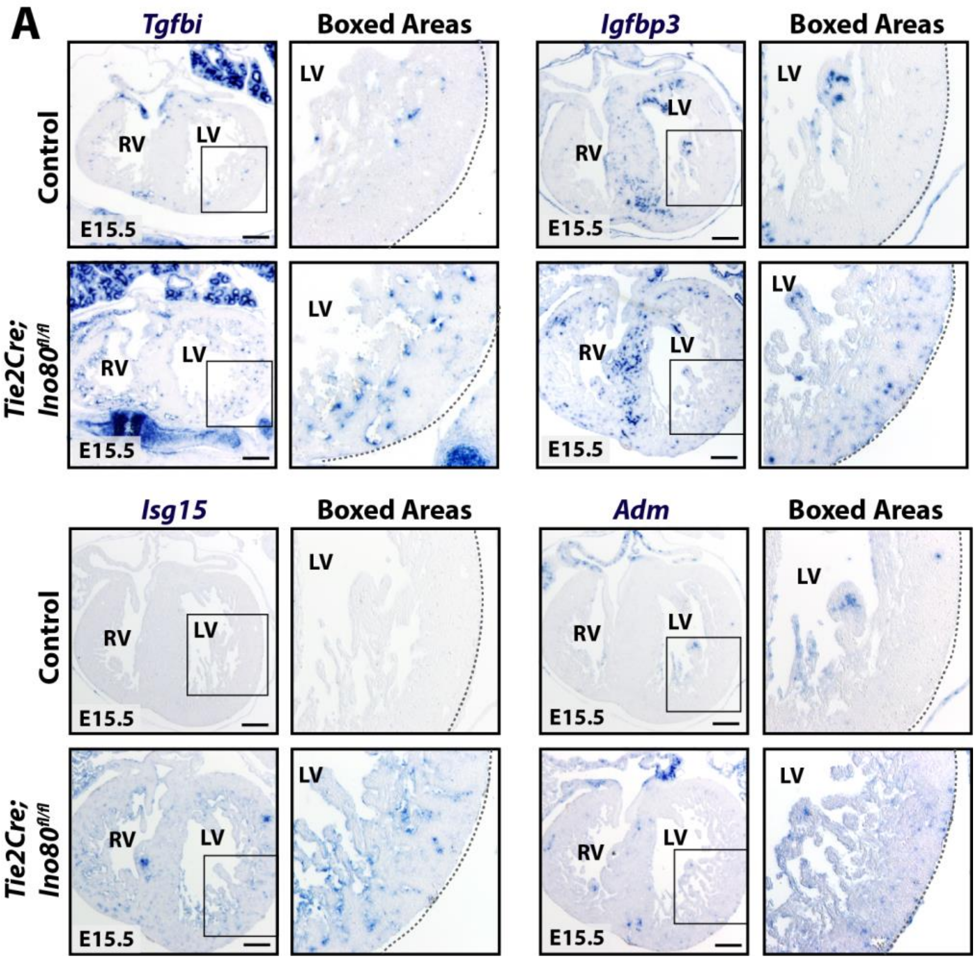


**s. Fig. 5 *In situ* validation of changes in indicated genes. (A)** *In situ* hybridization confirms the differences in gene expression of Tgfbi, Igfbp3, Isg15, and Adm in control and *Tie2Cre;Ino80*$^{fl/fl}$ hearts similarly as observed in the scRNA-seq data. Images are representative of the following number of replicates: control, n = 3 hearts; mutant, n = 3 hearts at E15.5. Scale bars: 25 µm.

### Supplementary Figure 6

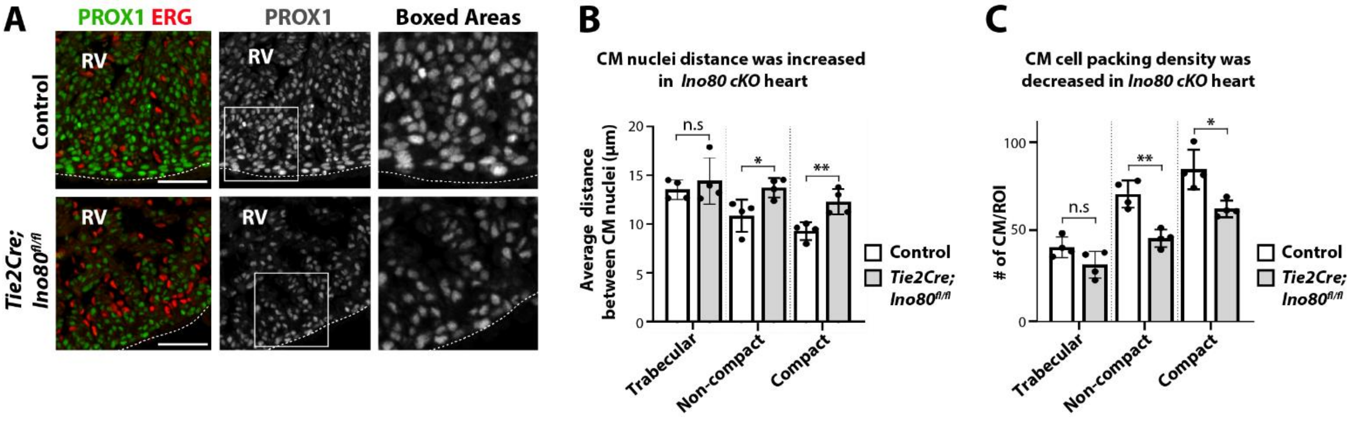


**s. Fig. 6. Nuclei distance and cell packing is changed in *Tie2Cre;Ino80$^{fl/fl}$* non-compacted hearts. (A)** Tissue sections stained for PROX1 (cardiomyocytes) and ERG (endocardium and coronary endothelium) demonstrate that mutants myocardium is less dense. Images are representative of the following number of replicates: control, n = 4 hearts; mutant, n = 4 hearts at E15.5. **(B)** Quantifying average distance between cardiomyocyte nuclei reveals an increase in non-compact and compact myocardium **(C)** Quantifying the total number of cardiomyocytes per region of interest (ROI) revealed reduced density in non-compact and compact myocardium. In bar graphs in **B-C**, each black dot represents the average value from >3 FOVs from one heart. Trabecular: trabecular myocardium; Non-compact: non-compact myocardium; Compact: compact myocardium. Scale bars: 25 µm (A). n.s: non-significant; *P < 0.05; **P < 0.01, evaluated by Student's t-test. Error bars are mean ± sd.

### Supplementary Tables

**Supplemental Table 1.** List of all differentially expressed genes in endocardial and endothelial cell subclusters

In a downloadable Excel spreadsheet titled ‘Supplemental Table 1’

**Supplemental Table 2.** List of all differentially expressed genes in each cardiomyocyte subclusters

In a downloadable Excel spreadsheet titled ‘Supplemental Table 2’

**Supplemental Table 3.** List of all differentially expressed genes in *Ino80* mutant endocardial and coronary vessel endothelial cells.

In a downloadable Excel spreadsheet titled ‘Supplemental Table 3’
